# BioNetComp: a Python package for biological network development and comparison

**DOI:** 10.1101/2021.04.14.439897

**Authors:** Lucas M. Carvalho

## Abstract

Due to the large generation of omics data on a large scale in the last few years, the extraction of information from biological data has become more complex and its integration or comparison as well. One of the ways to represent interactions of biological data is through networks, which summarize information on interactions between their nodes through edges. The comparison of two biological networks using network metrics, biological enrichment, and visualization consists of data that allows us to understand differences in the interactomes of contrasting conditions. We describe BioNetComp, a python package to compare two different interactomes through different metrics and data visualization without the need for a web platform or software, just by command-line. As a result, we present a comparison made between the interactomes generated from the differentially expressed genes at two different points during a typical bioethanol fermentation. BioNetComp is available at github.com/lmigueel/BioNetComp.

## Introduction

Networks are used to represent complex systems with relationships between their components. The networks are shown using a mathematical representation called a graph. A graph is made up of vertices and edges, and its application in several areas has already been observed, including in biological science (Zhang, P., & Itan, Y., 2019; Liu, C. et al., 2020; Mulder, N. J. et al., 2014). A biological network is generated using associations already known or predicted between components of a biological system, such as genes, proteins, metabolites, among others. With the availability of omics data on a large scale, both for model and non-model organisms, it was easier to study the interactions between biological systems, either through laboratory experiments or bioinformatics techniques. The comparative study of the interaction of these biological systems, represented by an interactome network, provides new insights into the understanding of systems biology.

There are several interaction databases of system components, such as MINT (Licatta, L et. al.,2012), DIP (Xenarios, I. et al., 2002), BIOGRID (Oughtred, R. et al., 2019), and STRING (Szklarczyk, D. et al., 2019). The latter has a database with more interactions and these are cured manually or are generated through bioinformatics techniques, such as coexpression and text mining (Harrington, E. D., Jensen, L. J., & Bork, P., 2008). Also, STRING has a web application that is limited to receiving a list of up to 2000 genes or proteins and generating an interactome. This generated network can be opened in software that performs network analysis, such as Cytoscape (Shannon, P. et al., 2003). It has modules to enable the execution of a general analysis of the network, such as basic metrics and enrichments. The comparison between two networks can also be performed by Cytoscape, through the Dynet package (Goenawan, I. H., Bryan, K., & Lynn, D. J., 2016), but it has a difficulty in dealing with extremely dense networks and generating reports. There is also a web application, called NetConfer (Nagpal, S. et al., 2020), which performs the comparison of several networks at the same time using already known metrics, but networks are not generated automatically.

In this article, we present BioNetComp, a python package that compares two biological networks and their metrics. Also, BioNetComp can be executed on the command line, and it generates reports that can be used in various software that interpret networks.

## Material and Methods

### Dataset

We assessed the performance of BioNetComp through a publicly available dataset (Carvalho-Netto, O. V. *et al.*, 2015) that allows comparing the interactomes generated over a typical industrial fermentation of bioethanol, also called 1G fermentation. The complete dataset of RNA-Seq reads can be accessed in SRA accession SRA057038.

### RNA-Seq analysis

For each RNA-seq library, reads were aligned to S. cerevisiae S288c genes (www.yeastgenome.org) using the kallisto v1.0.7 software (Bray, N. L. et al., 2016). The differential expression analysis was performed using the DESeq2 v1.30.1 package (Love, M.I. et al., 2014) which has an option to deal with datasets without biological replicates.

### Network development

Interactions were requested in the STRING v11 database through an API. Networks and metrics were generated using the networkX v2.5.1 package (Hagberg, A. A. et al, 2008).

### Stress tests

Stress tests were implemented in Python 3.8.5 and executed in a Ubuntu Server on Intel(R) Xeon(R) CPU E5-2420 (2.20 GHz), 48 GB of RAM. Each instance was executed 10 times and the mean was used in further analysis. The use of memory throughout the BioNetComp execution was verified by the memory profiler package (https://github.com/pythonprofilers/memory_profiler).

### Data availability

The source code for BioNetComp is available in an online repository (github.com/lmigueel/BioNetComp).

## Workflow

BioNetComp contains a flowchart designed to provide a structured comparative approach between two biological networks through the STRING database, as well as metrics, comparative reports and network visualizations.

From the entry of two lists of proteins or genes and the taxid of the organism under study, we execute the pipeline described in Figure 1. First, a request is generated in the STRING database, limited to 2000 proteins/genes, to return the interactions described in each biological network. Subsequently, from the interaction pool, the network is generated through the NetworkX package. Finally, comparative reports and a final network are generated, and colored from the presence or absence between the networks. For more detailed information, access the repository: github.com/lmigueel/BioNetComp.

**Figure 1.**
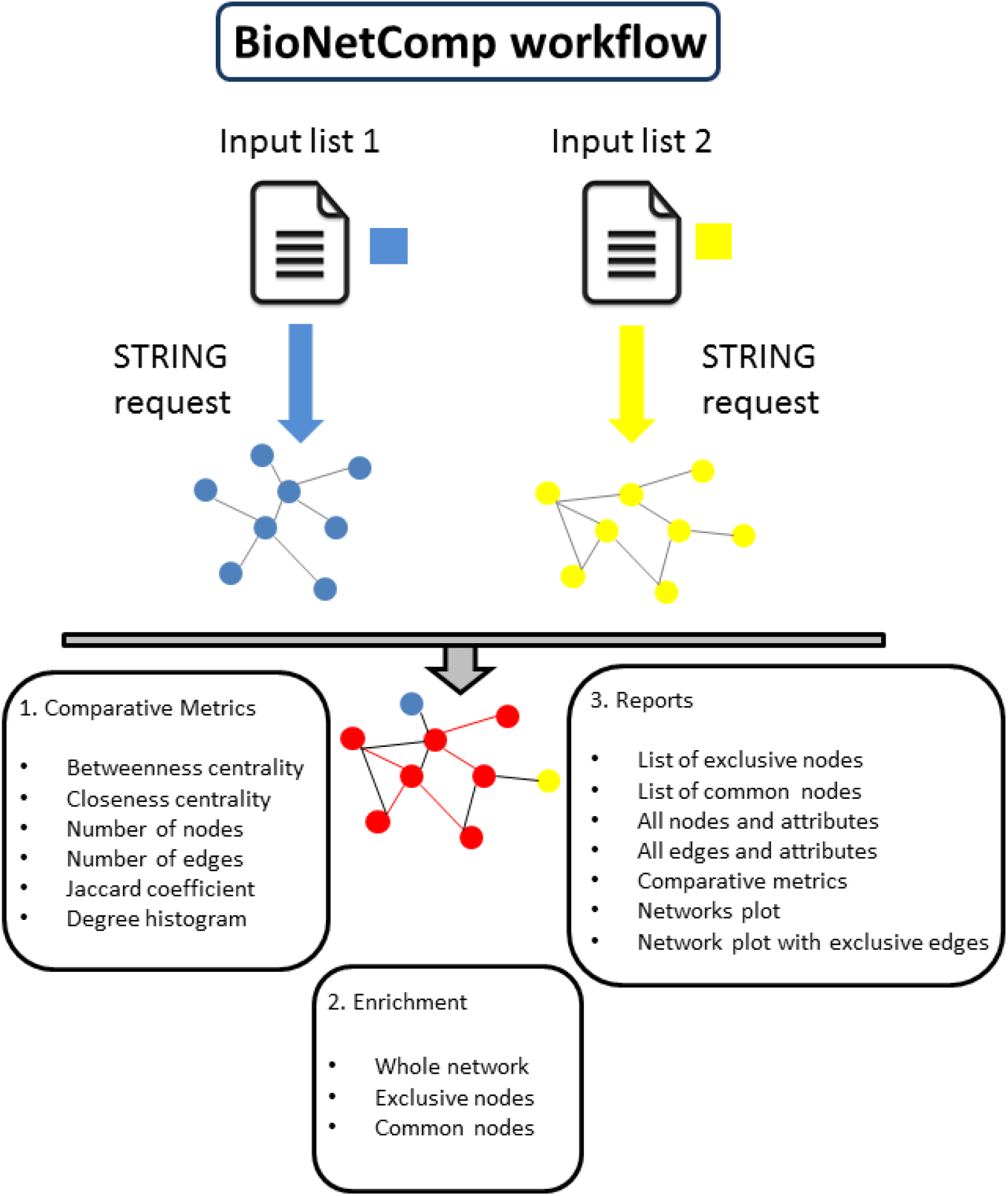
BioNetComp workflow. From two lists of genes or proteins, a request in the STRING database is carried out and the interactions are stored. From this, biological networks are generated. The comparison is made using several metrics and includes comparative charts. Biological enrichment of exclusive and similar nodes is also carried out. Finally, a final report is generated, including the graphs of each network and a final network, containing all the information about exclusivity and similarity between nodes and edges.

The reports provided by BioNetComp from two lists of genes or proteins are:

1. A text file containing the list of nodes and total edges, differentiated by color and presence and absence in the network;
2. A text file containing exclusive nodes of each network and those in common;
3. Exclusive networks and a final network plot, containing comparative information;
4. Network plot generated only by exclusive edges of each biological network;
5. Comparative graphics of the number of nodes and edges;
6. Exclusive comparison charts, such as the betweenness and closeness centrality boxplots;
7. Degree histogram chart and its boxplot for each network;
8. Enrichment of the entire network, but also exclusive and common nodes.
9. Betweenness and closeness centrality gene ranking;
10. Jaccard coefficient between networks applied to nodes and edges for dissimilarity observations.

### Network development

The first step of BioNetComp is to make requests in the STRING database through its API, which enables you to get the data without using the graphical user interface. These requests are stored in a data frame and an edge hash, which stores the interactions found. The requests depend on taxid number, which is a required option in our package (*--taxid*). If the number of proteins or genes exceeds 2000, a warning message will be displayed.

The interactions found in the STRING database are transformed into a data frame that can be read by the NetworkX package through the *from_pandas_edgelist()* module. With the network graphs, all comparative metrics can be generated. The user can change the threshold value for STRING interaction score using the *-- threshold* option, which has a default value of 0.40. For details about STRING interaction score calculation access http://version10.string-db.org/help/faq/.

The module present in BioNetComp for Network development is called *network_development()*.

### Comparative Metrics

With the networks, we can calculate their basic metrics and compare them. The first step is to compare the total nodes and edges of each network. A bar chart is generated.

Some other network metrics are also generated and compared. The first is the closeness centrality, which indicates how close a node is to all others in the network. The second metric is the betweenness centrality, which measures the importance of each node in passing information, that is, it deals with the identification of hub genes. Finally, the histogram degree for each network is produced. The comparative boxplot for these metrics is also generated by BioNetComp.

The final report for each network contains (i) number of vertices; (ii) the number of edges; (iii) whether the network is connected or not; (iv) average path length and average diameter if the network is connected; (v) network diameter of the largest component; (vi) average clustering; (vii) node with max closeness centrality; and (viii) node with max betweenness centrality.

### Network visualization

In addition to the comparative charts for each metric mentioned in the previous section, BioNetComp generates the network plots. The nodes that have the highest betweenness centrality value are highlighted with a larger size due to their importance in the network. The graph contains a color scale, indicating the value of the degree, therefore, nodes with a greater tendency to be a hub are also highlighted. Thus, through this visualization technique, we highlight essential nodes in the network. We use the spring layout of the NetworkX package, as it can better sample the nodes through the optimal distance between nodes.

The module present in BioNetComp for Comparative metrics and Network visualization is called *comparative_metrics()*.

### Network Enrichment

The network enrichment is performed through requests using the STRING API. The user can change the FDR cut-off value for enrichment using the *--fdr* option, which has a default value of 0.05.

The module present in BioNetComp for network enrichment is called network_*enrichment()*.

### Network Comparison

The comparison between the networks is carried out in a simple, but practical and objective way. First, reports are generated containing the exclusive nodes of each network and the nodes in common. Also, a report of the edges of the final network is generated, containing the edges that are shared by both networks and the exclusive ones. The attributes of each edge are also reported. The enrichment of each node group is also reported, so we can identify the unique and common biological processes.

With the basic comparison metrics generated, we calculate the Jaccard coefficient for nodes and edges. This coefficient measures the similarity between the observed metrics. At the end of this step, a bar chart is generated containing the value of the Jaccard coefficient for the nodes and edges.

The final plot of the network contains information that allows a clear visual comparison. The nodes that are common to both networks, exclusive of the first network (*--in1*) and exclusive of the second network (*--in2*) will be colored by red, blue and yellow, respectively. Also, the degree of each node delimits the size of the node in the final network plot, allowing visualization of the hub nodes.

The module present in BioNetComp for network comparison is called *compare_networks()*.

The reports names always take into account the data entry. The label ‘network1’ will be assigned to the list of genes or proteins in *--in1* option, while the label ‘network2’ will be assigned to the list present in the *--in2* option. All reports will be generated within the folder present in the *--output_folder* option.

## Results and discussion

### Case of study: Understanding differences between differentially expressed genes during bioethanol fermentation

Carvalho-Netto, O. V. *et al.* performed an RNA-Seq analysis during typical bioethanol fermentation. The collected points were 1h, 4h, 7h, 10h, 12h, and 15h during the fermentation. Then, the paired differential expression of all points was performed based on the first time-point (1h). The results of measured metabolites presented by the authors show that C6 sugar (glucose) is consumed within 12 hours of fermentation, with the highest consumption rate between 7h and 10h. Also, throughout the fermentation, glycerol is produced. We will perform the comparison of the interactome from differentially expressed genes of 4h *versus* 1h and 12h *versus* 1hr comparisons. The chosen points are contrasting both in the production of ethanol and the consumption of glucose, which is more accentuated at the beginning of fermentation.

The first network (network1) was generated from the 306 differential genes between 4h and 1h. The second network (network2), on the other hand, was generated from 1559 differential genes between 12h and 1hr. We observed a difference between the number of network final nodes (279 vs 1559) and edges (2778 vs 32289). We also note that network1 is disconnected and network2 is connected, which ends up causing a difference in betweenness centrality (Fig. 2A), closeness centrality (Fig. 2B), and degree distribution (Fig. 2C-D). The second group of differential genes (network2) generates a connected network, and this causes the value of closeness centrality to increase, as shown in Fig. 2B, and higher values of the degree distribution (Fig. 2D). We also note that there is a very connected group in the first group of differential genes (network1), which can be seen highlighted in yellow in Fig. 2E. This perturbation causes two peaks in the degree histogram (Fig. 2C) and ends up increasing the value of betweenness centrality (Fig. 2A). The RPS31 gene is the node with the highest value of closeness centrality in both networks, and its main role is to act on ribosomal biogenesis and assembly. A dense final network is shown in Figure 3.

**Figure 2.**
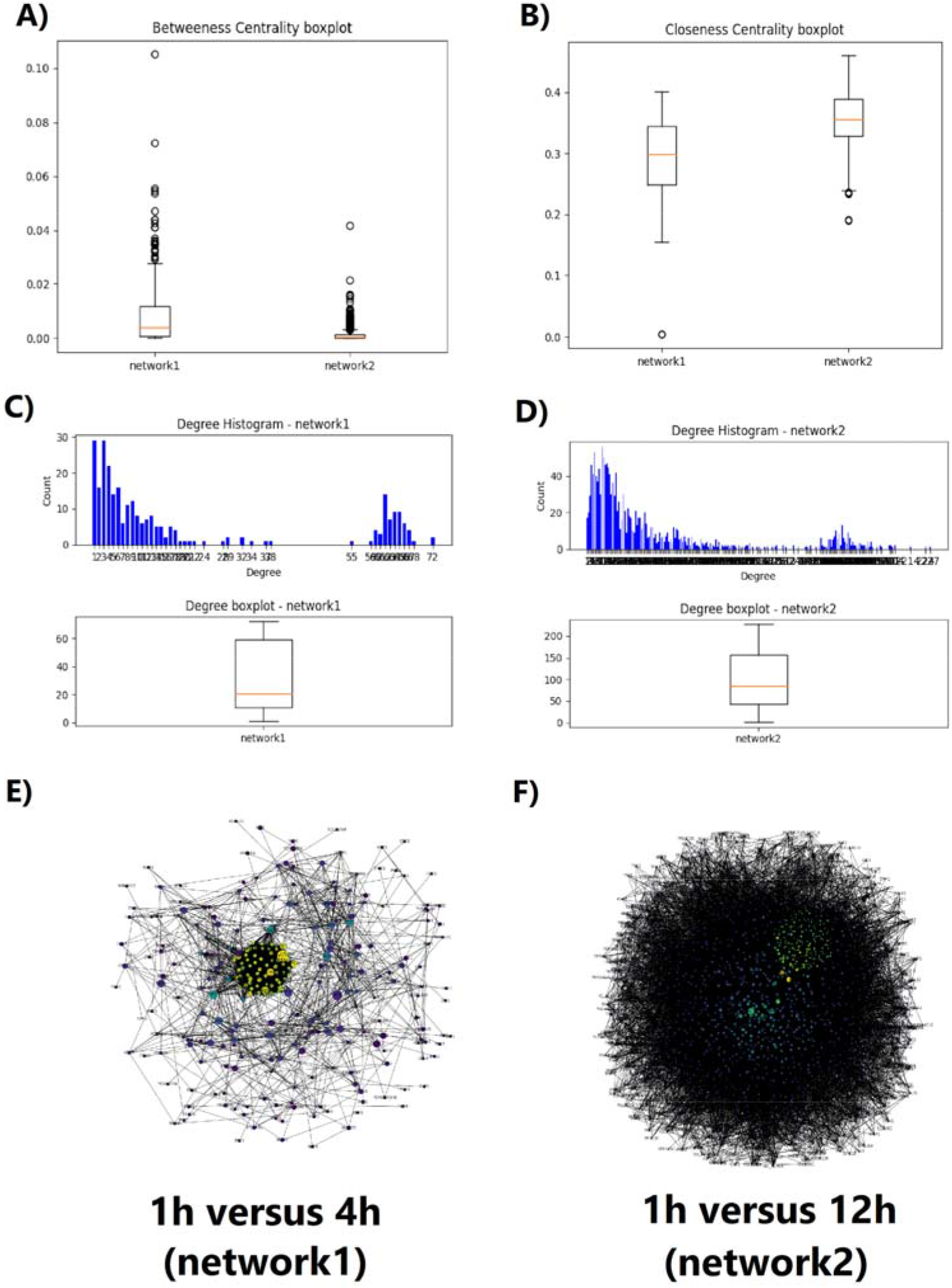
Some of the final results obtained by BionetComp in the comparison of the interactome generated through the differentially expressed genes of the points 1h versus 4h and 1h versus 12h in the typical bioethanol fermentation described by Carvalho-Netto et. al., 2015. (A) Comparative boxplot of betweenness centrality. (B) comparative boxplot of closeness centrality. (C) Degree histogram and boxplot for the first group of genes. (D) Degree histogram and boxplot for the second group of genes. (E) The final network plot for the first group of genes. (F) The final network plot for the second group of genes.

**Figure 3.**
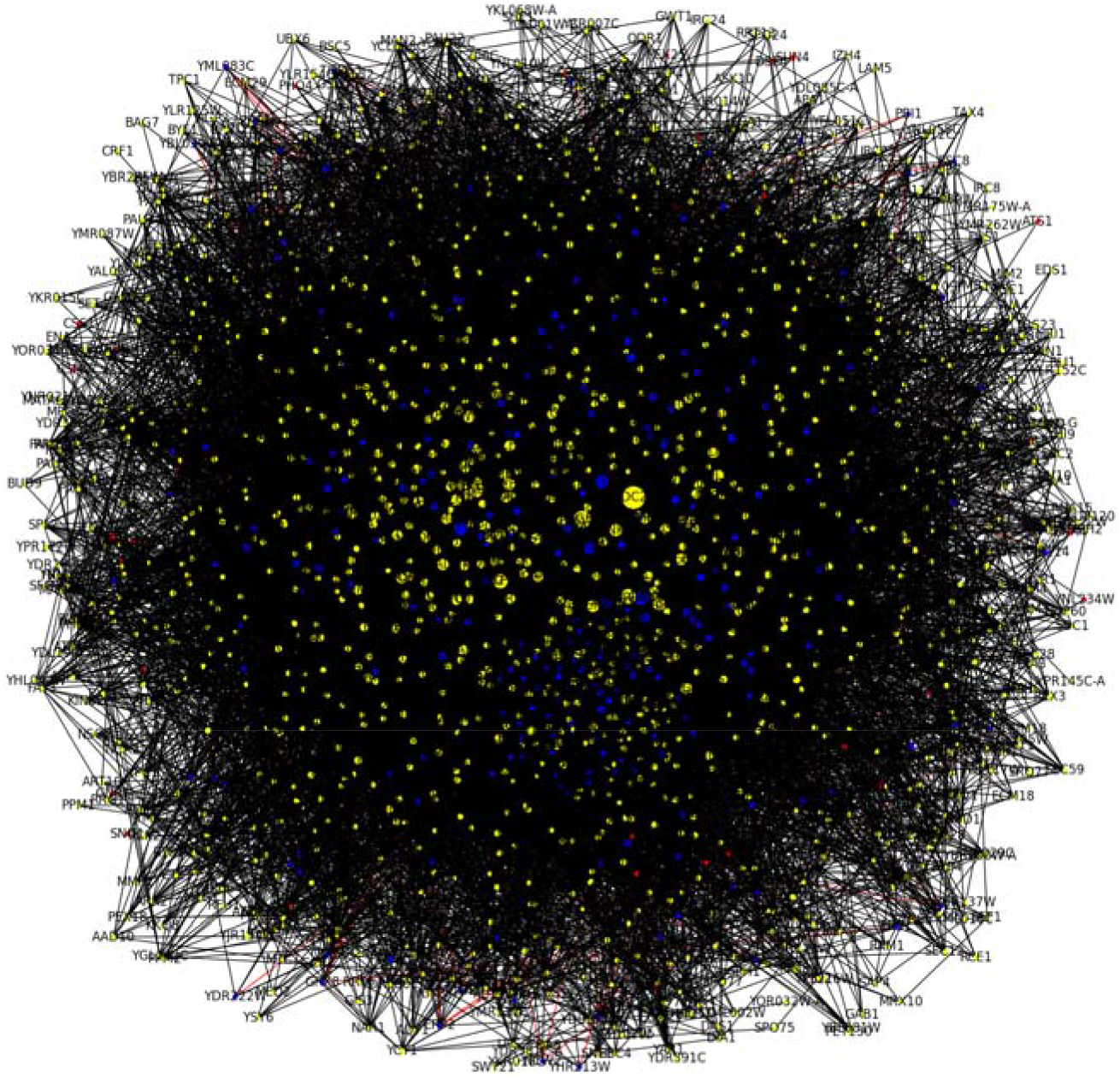
Final network plotted by BioNetComp. Nodes in red, blue, and yellow represent the intersection nodes, exclusive nodes of network1, and exclusive nodes of network2, respectively. This network is generated by the report files edge_report.txt e node_report.txt.

Regarding the results of the enrichment of the nodes in common between the networks, we note that there are many enriched GO processes (p-value <= 0.05) involving in ribosomal roles (GO: 0042255, GO: 0042254, GO: 0042273, GO: 0000027, and GO: 0042274), related to processes of transcription and cellular organization (GO: 0006405, GO: 0010467, GO: 0051029 and GO: 0031505) and metabolic processes and stress (GO: 0034599 and GO: 0006083). The exclusive nodes of network1 have enriched GO processes (p-value <= 0.05) related to glutamate metabolism (GO: 0006536, GO: 0009084 and GO: 0006537), glycolysis, and pentose phosphate (GO: 0019682, GO: 0019323, GO: 0019693), repair (GO: 0000730, GO: 0090735 and GO: 0000725) and mainly in metabolic processes involving oxidative stress and NAD / NADH (GO: 0034354, GO: 0034627, GO: 0019674, GO: 0006739 and GO: 0051252). These results from network1 show that yeast, at the beginning of fermentation, already suffers from a very large redox imbalance in the industry, which is also being corrected by glutamate processes but has important metabolic processes to produce ethanol. The exclusive nodes of network2 have many enriched GO processes (p-value <= 0.05) related to cellular respiration (GO: 0022904, GO: 0045333, GO: 0042775, GO: 0009060, GO: 0006122, and GO: 0006121), and autophagy and starvation (GO: 0000422, GO: 0009267, GO: 1903008, GO: 0000422, GO: 0061912). This shows that the yeast, at the 12h fermentative point, tries to overcome the redox imbalance by activating the mitochondrial process, but enters a state of starvation due to the lack of sugar.

Pathway enrichment analysis over common nodes shows metabolic pathways related to bioethanol fermentation, such as Glyoxylate and dicarboxylate metabolism, Glycolysis/Gluconeogenesis, beta-Alanine metabolism, Fatty acid elongation, and Ribosome. However, network2 has metabolic processes related to the end of fermentation and starving beginning, such as Autophagy, Citrate cycle (TCA cycle), Meiosis, DNA replication, Oxidative phosphorylation, and Starch and sucrose metabolism. Network1 has not any exclusive pathway enriched.

All results generated by the BioNetComp package for this comparison can be found in the Supplementary Material A.

### Stress tests

To test the limits of BioNetComp and evaluate its behavior, a stress test was carried out for a different number of genes/proteins in each network. Two random networks were generated from *Saccharomyces cerevisiae* genes. The total number of genes present in each biological network varies from 200 to 2000. The time spent was calculated during the execution of the four essential steps of BioNetComp: (i) Step1: STRING API request; (ii) Step2: Network development, Comparative metrics, and Network Visualization; (iii) Step3: Network enrichment; (iv) Step4: Network comparison. The results are summarised in Figure 4.

**Figure 4.**
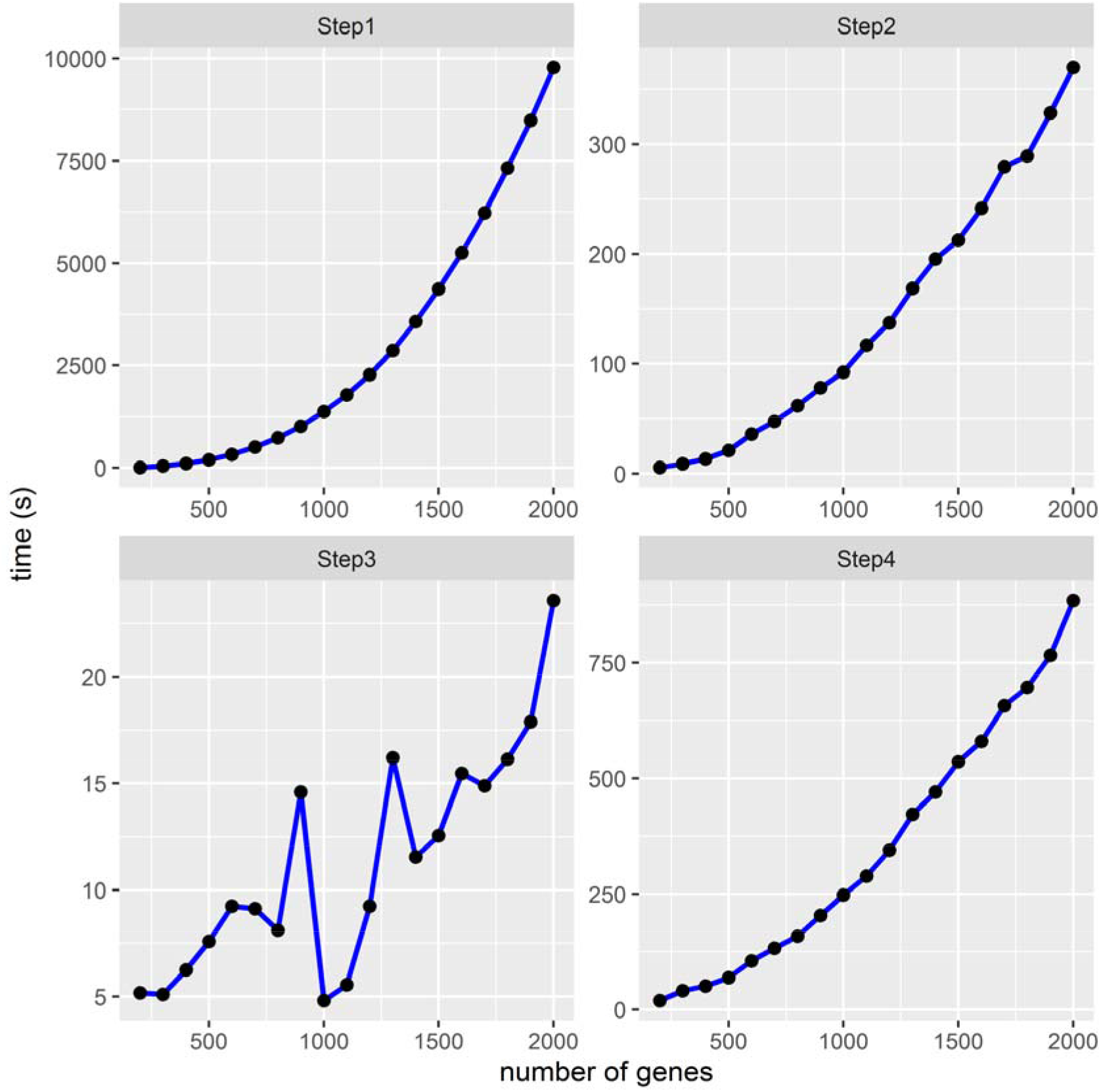
Stress test during BioNetComp execution. The x-axis represents the total number of genes in each biological network selected randomly from the *S. cerevisiae* genes, which varies between 200 and 2000 nodes. The y-axis shows the time spent on each processing step. The steps are: (i) Step1: STRING API request; (ii) Step2: Network development, Comparative metrics, and Network Visualization; (iii) Step3: Network enrichment; (iv) Step4: Network comparison.

The use of memory throughout the execution of BioNetComp was verified by the memory profiler package. We note that the highest memory usage (in Mb) occurred when both networks have 2000 nodes and was limited to the total usage of 500 Mb. All memory usage results are available in the Supplementary Material B.

## Conclusions

Given the established importance of interpreting biological information through networks, there is also an inherent need for tools that can compare and visualize them more objectively and clearly. The general intention behind the development of BioNetComp is to generate results quickly, easy and not dependent on web platforms or software. Despite the limit for very dense networks, the metrics generated in the reports and the comparative graphs are essential to extract conclusions from the interactomes. The command-line use of BioNetComp is expected to facilitate the automation of processes that need to extract information from biological networks.

## Ethics declarations

Not applicable

