## Supplementary figures and images for "BioNetComp: a Python package for biological network development and comparison"

### betweenness_centrality_boxplot.png

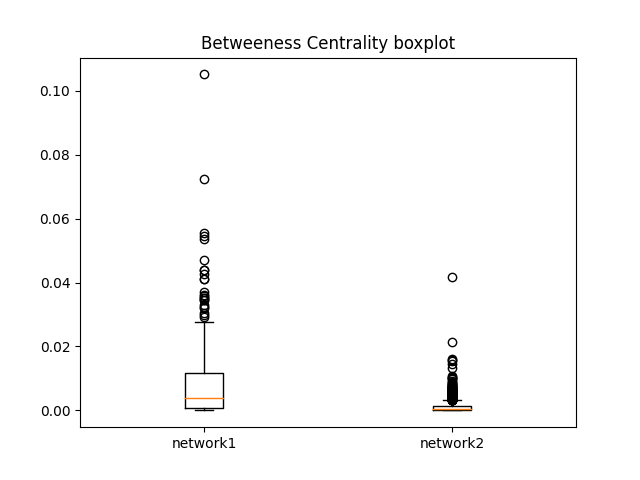

### closeness_centrality_boxplot.png

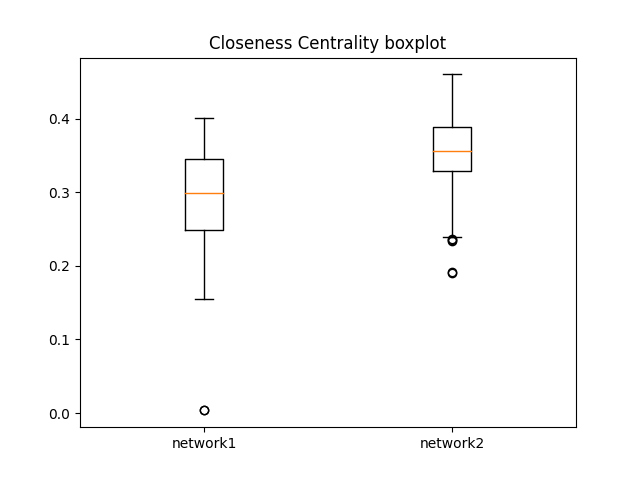

### final_network_plot.png

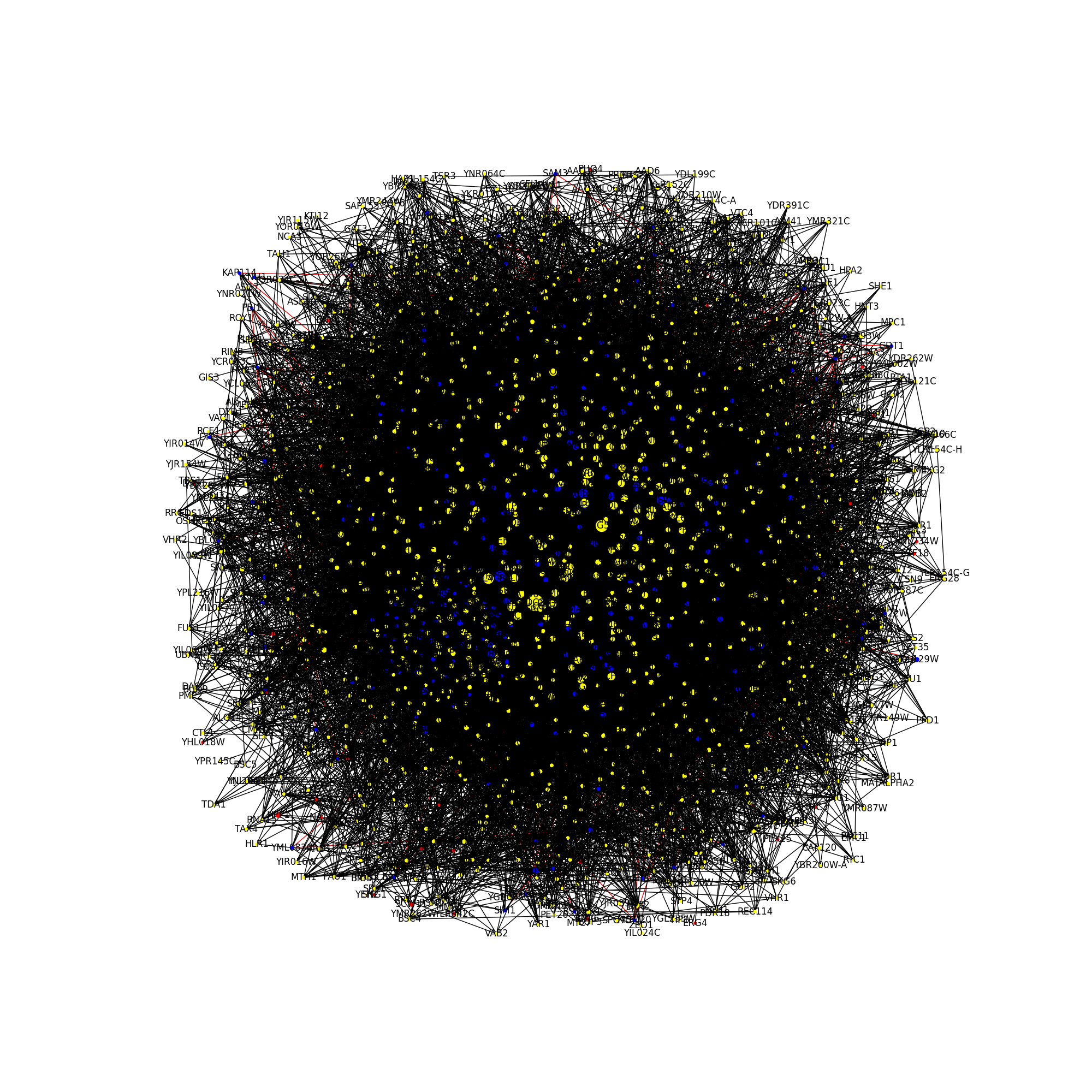

### jaccard_coefficient_nodes_and_edges.png

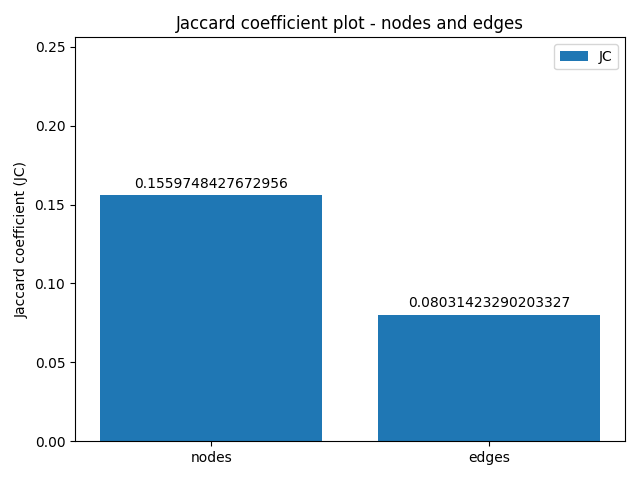

### network1_degree_hist.png

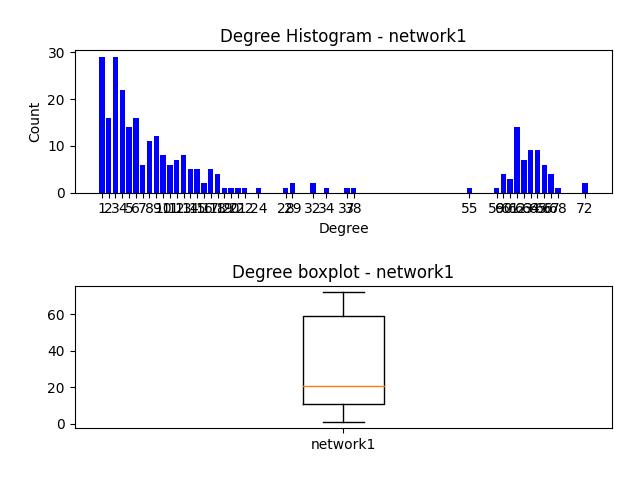

### network1_plot.png

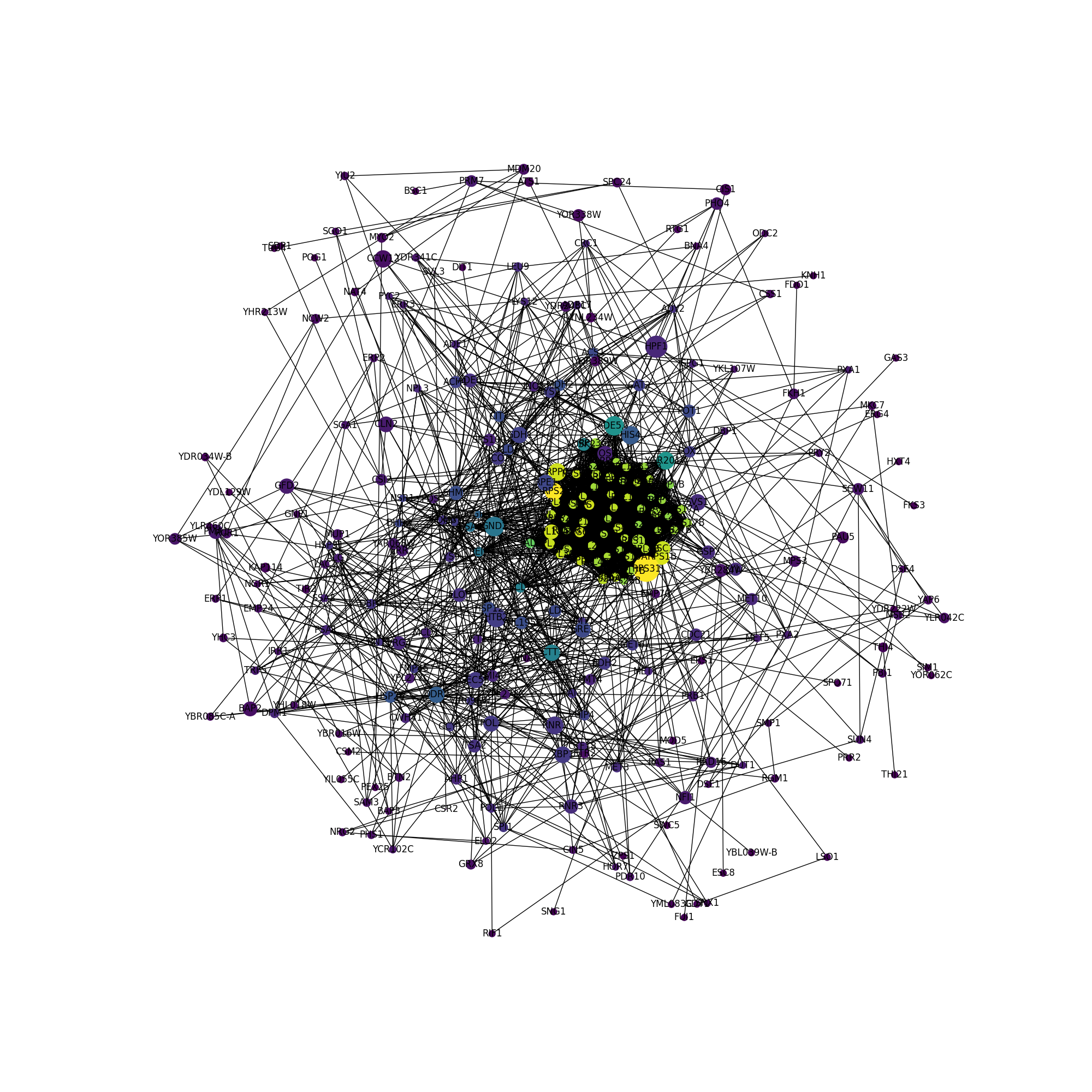

### network2_degree_hist.png

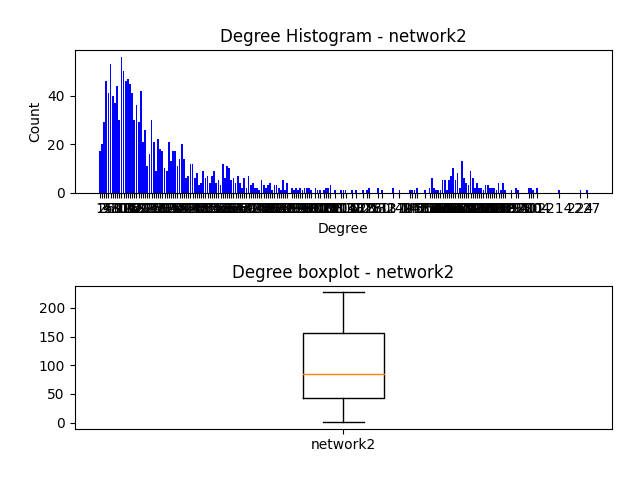

### network2_plot.png

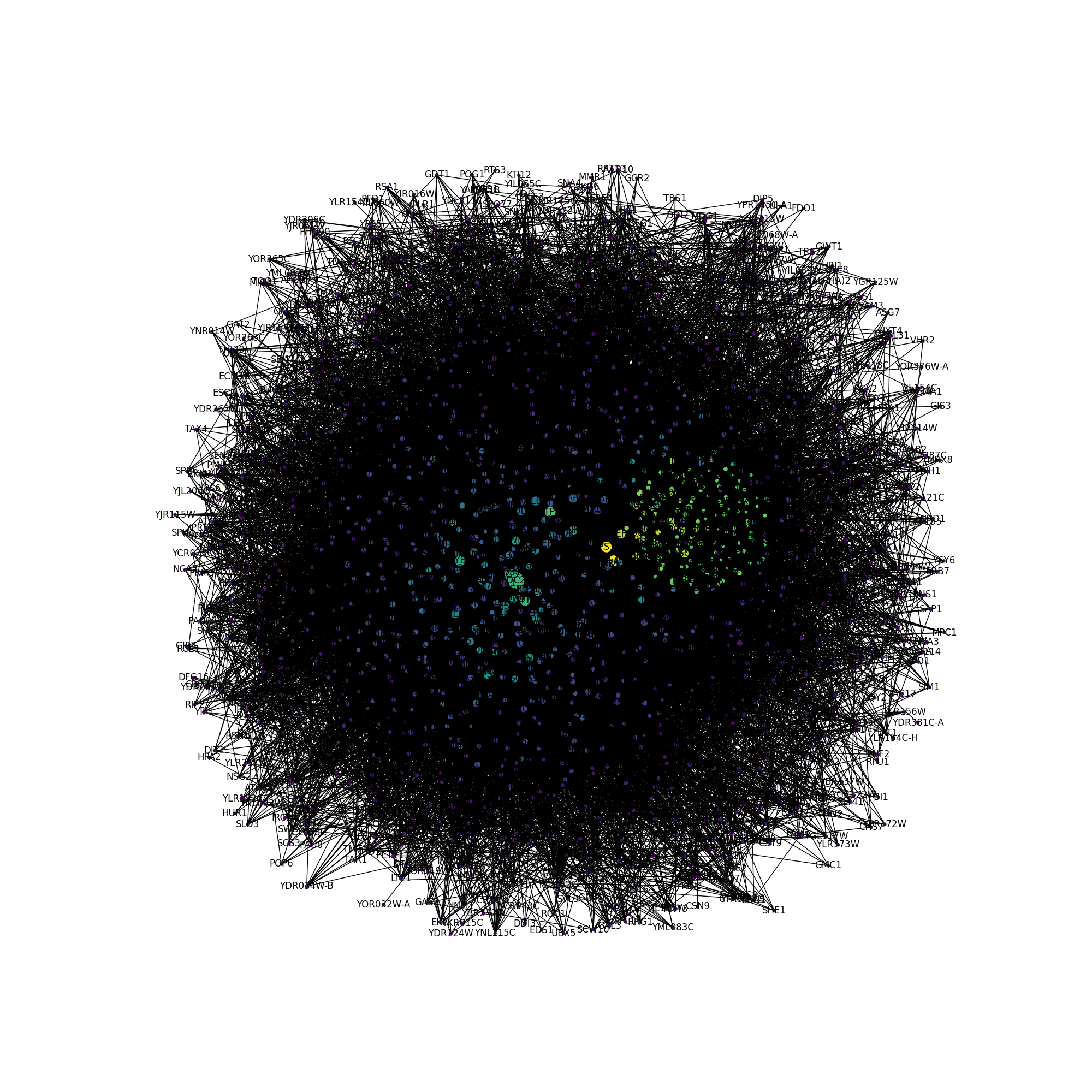

### nodes_edges.png

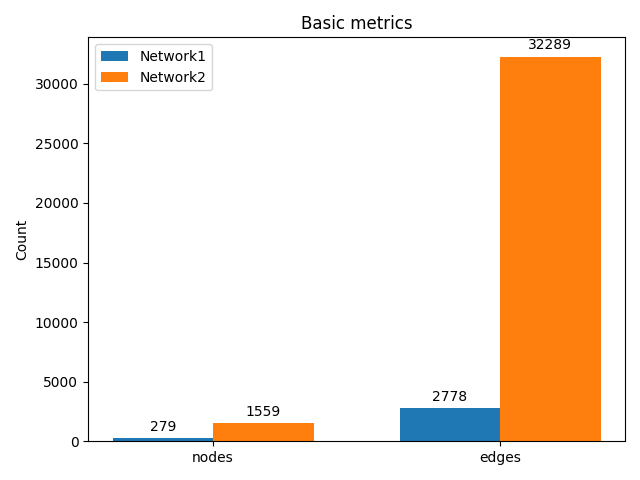

### only_network1_edges.png

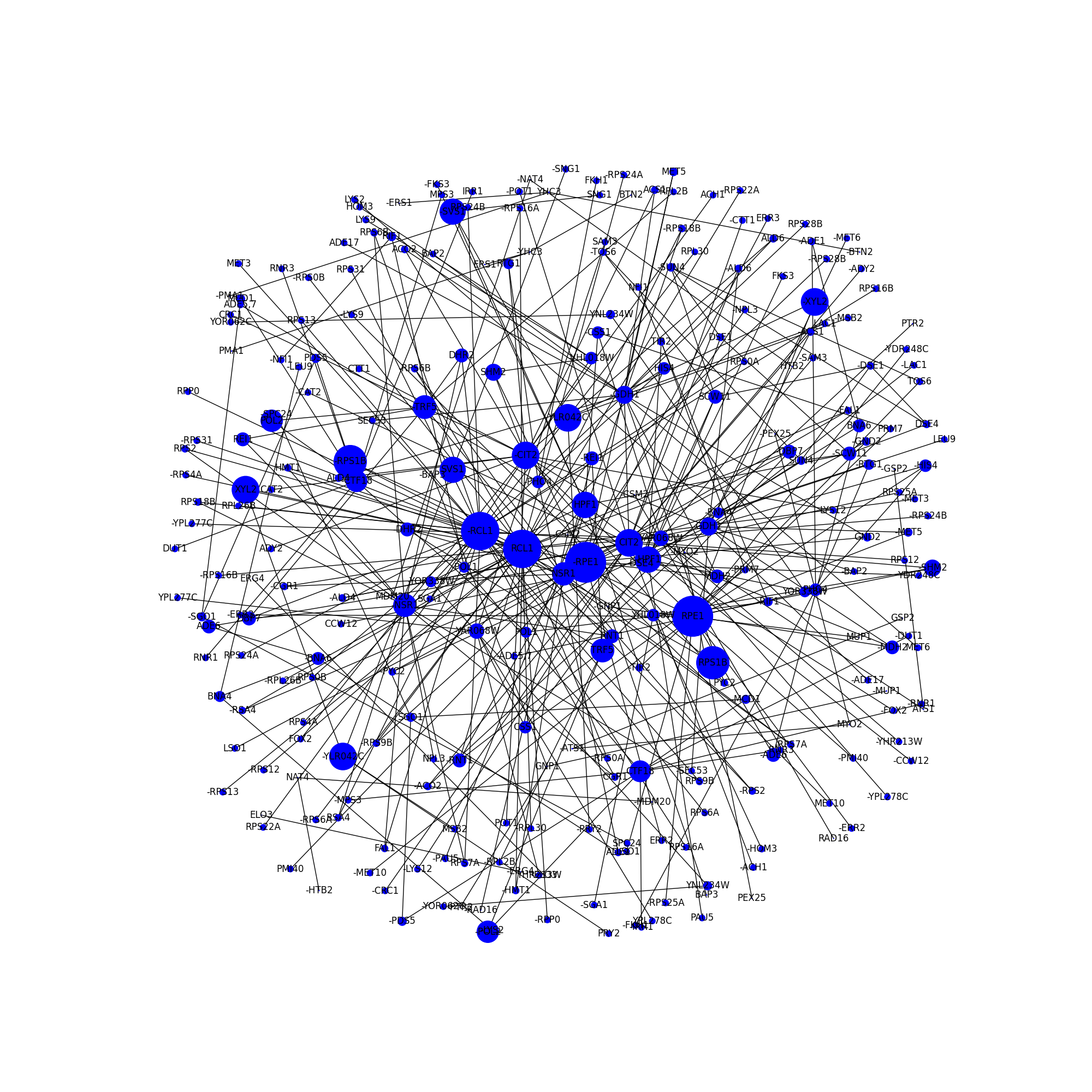

### only_network2_edges.png

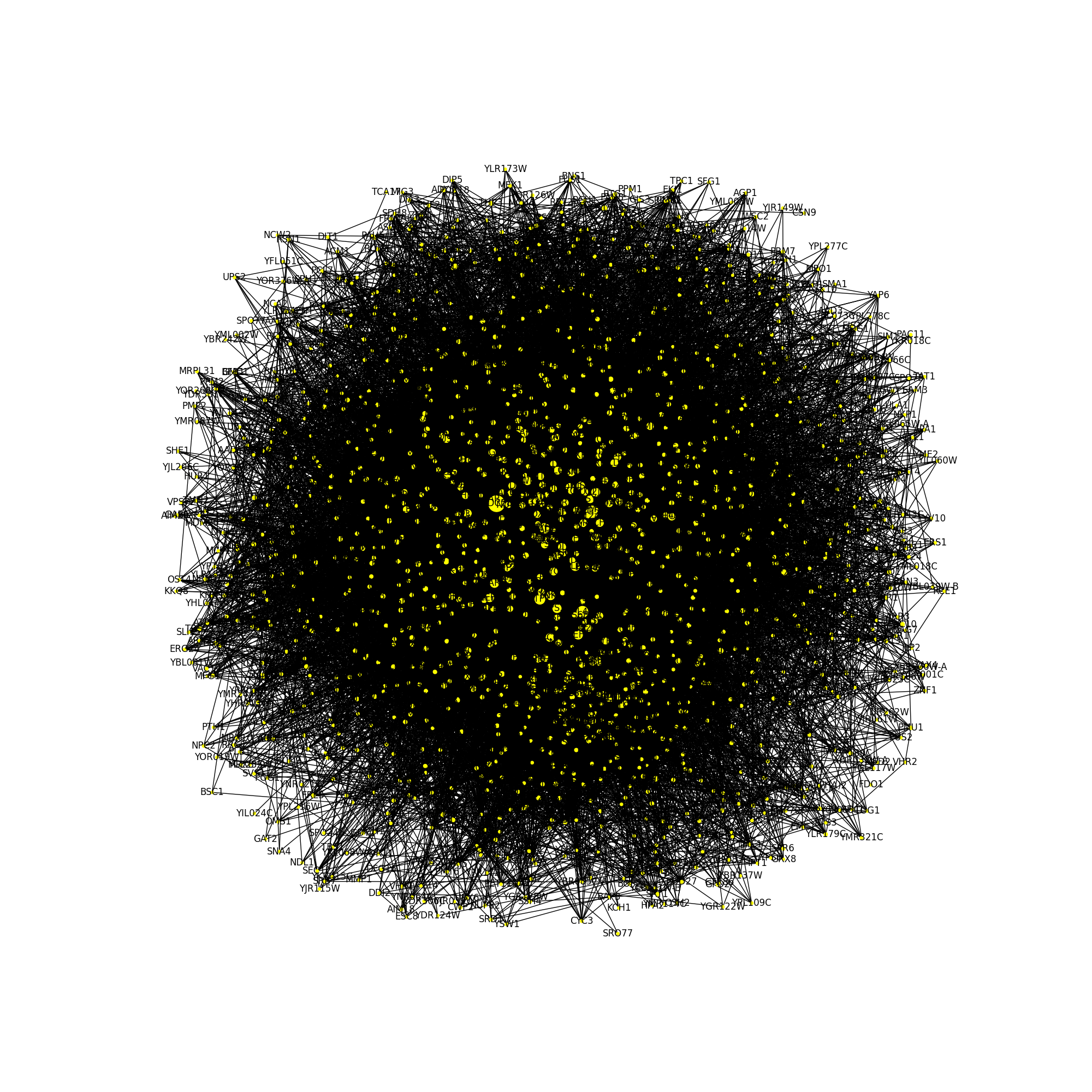

### plot_n200.png

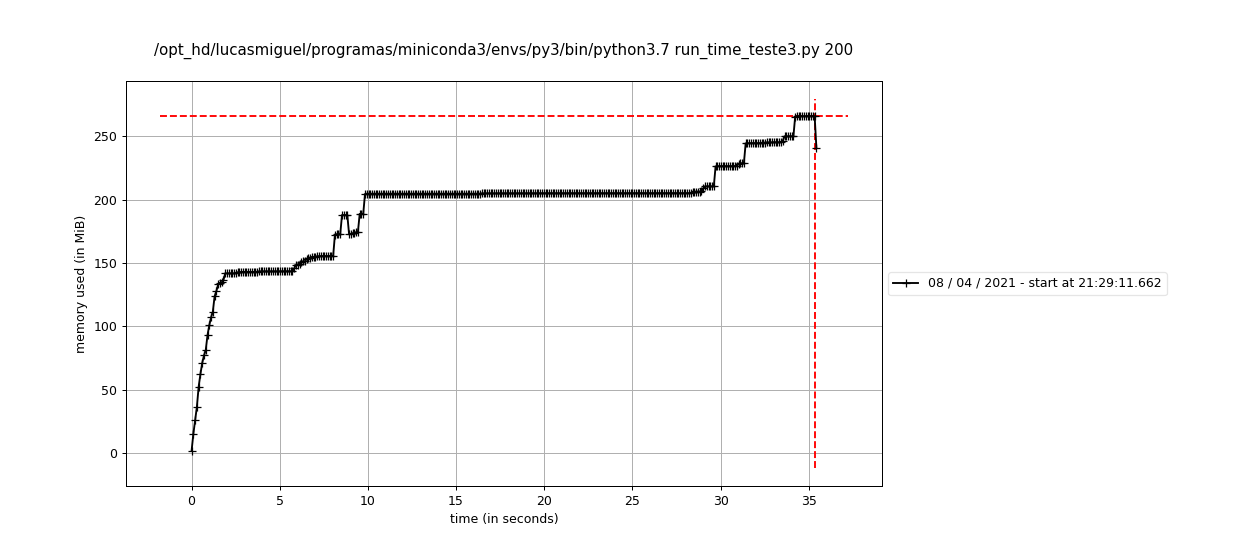

### plot_n300.png

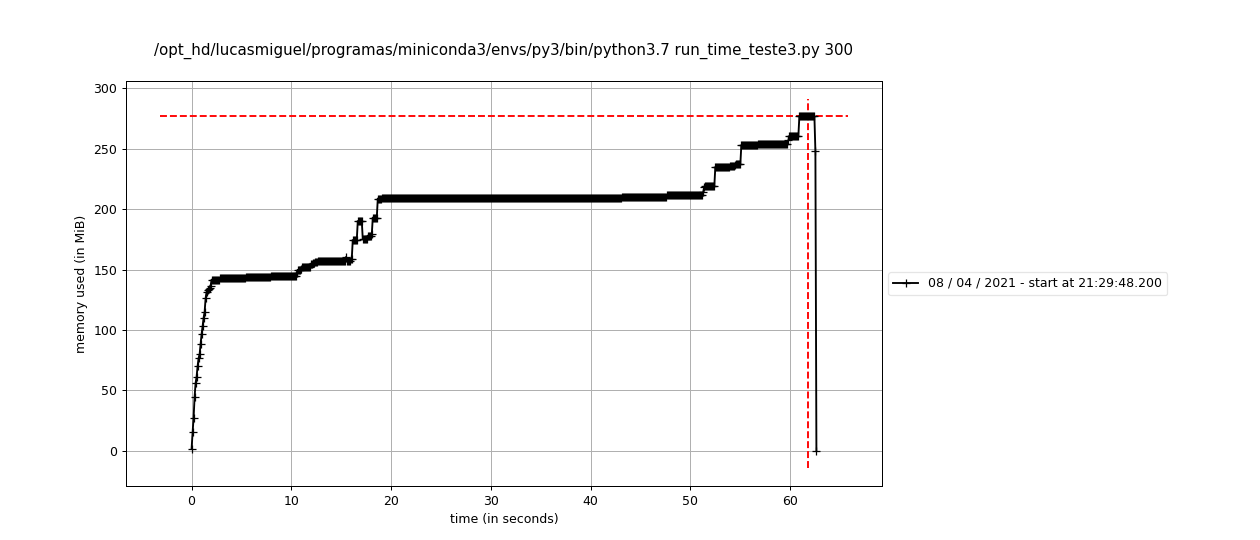

### plot_n400.png

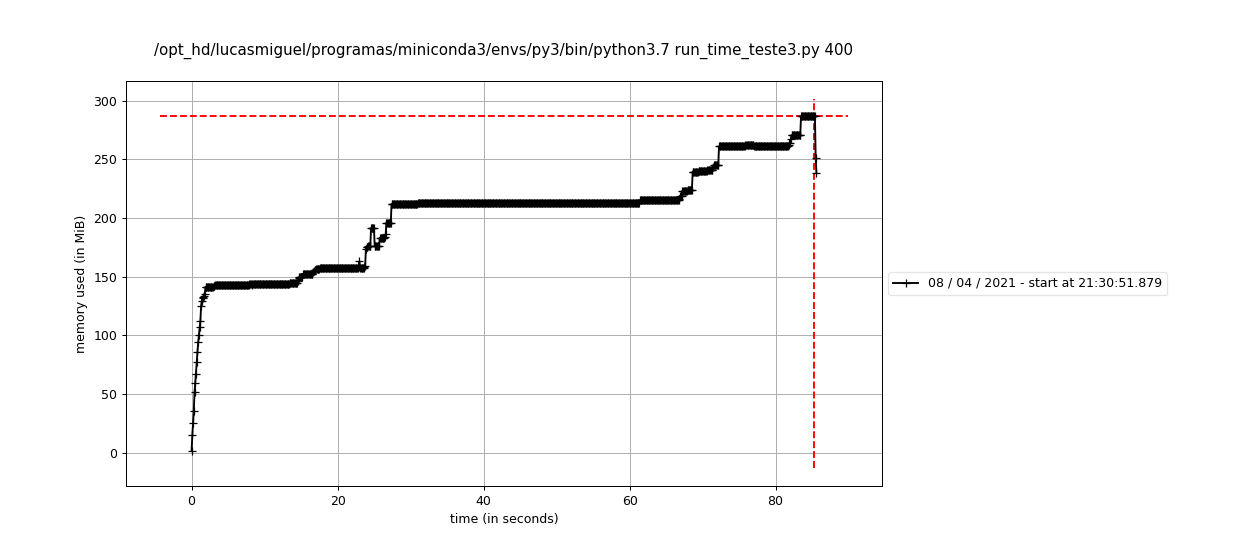

### plot_n500.png

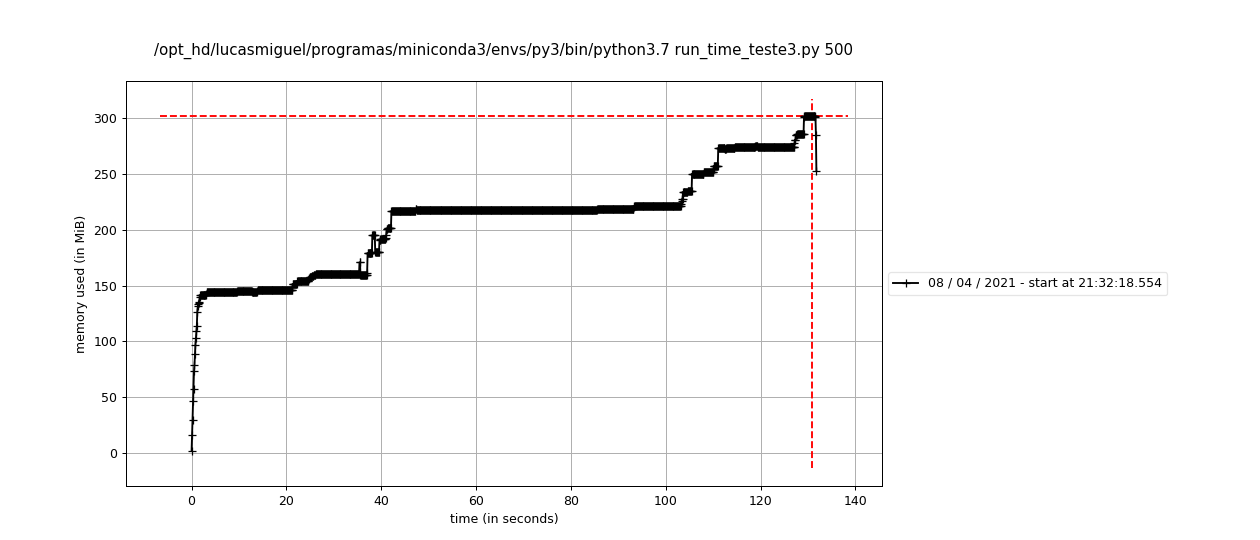

### plot_n600.png

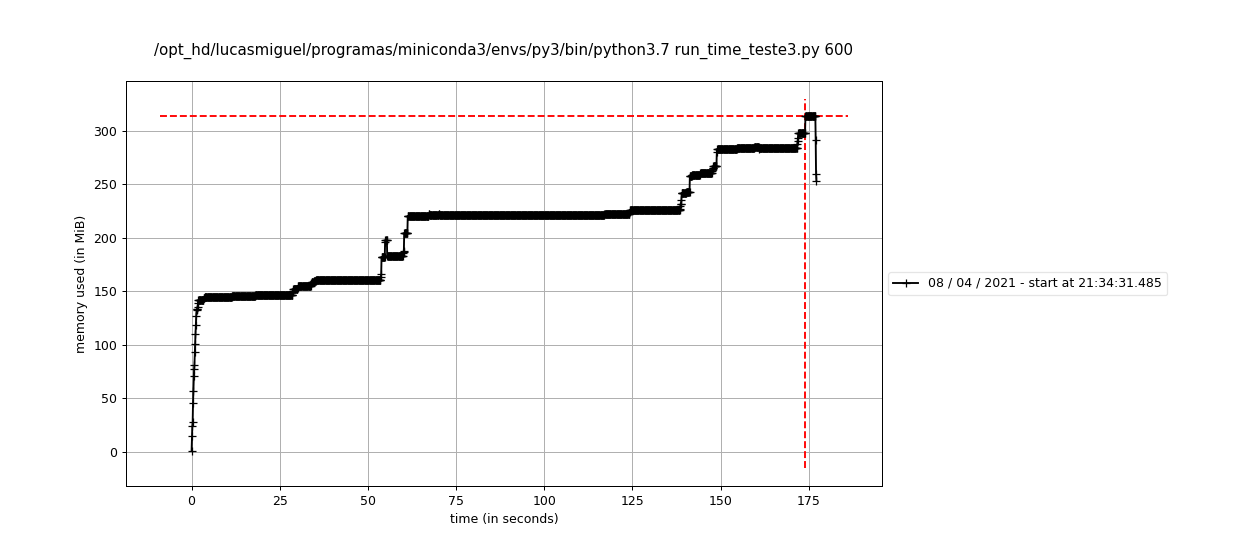

### plot_n700.png

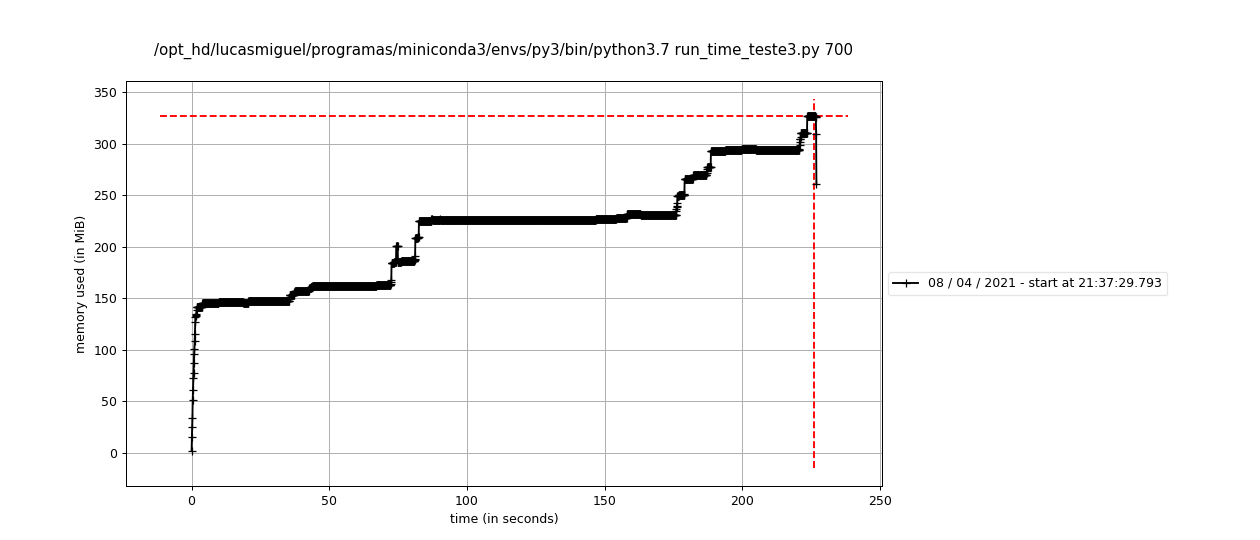

### plot_n800.png

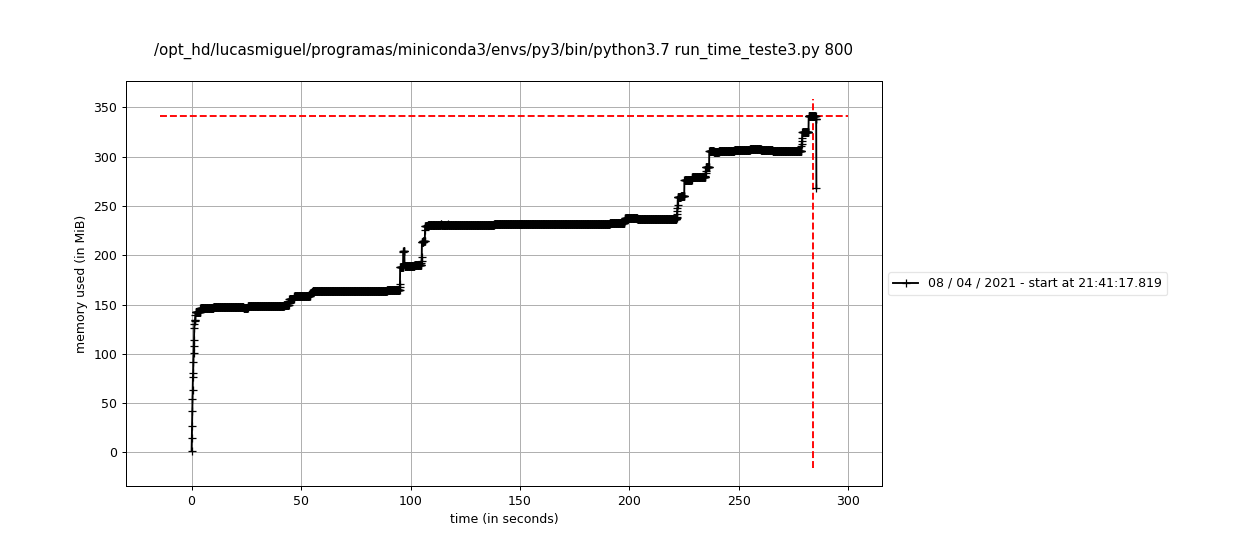

### plot_n900.png

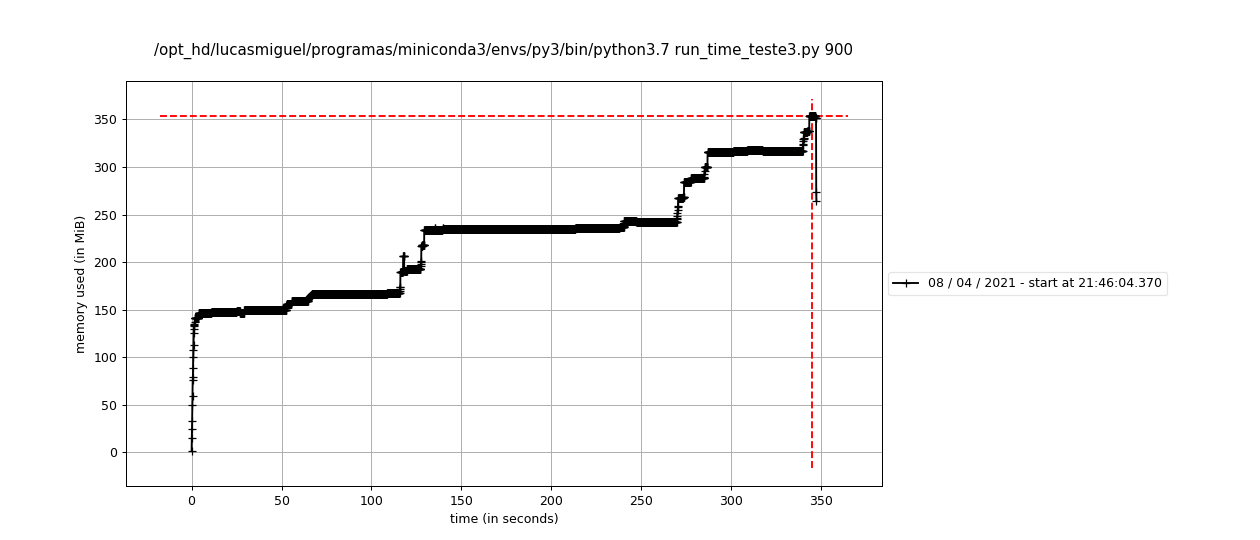

### plot_n1000.png

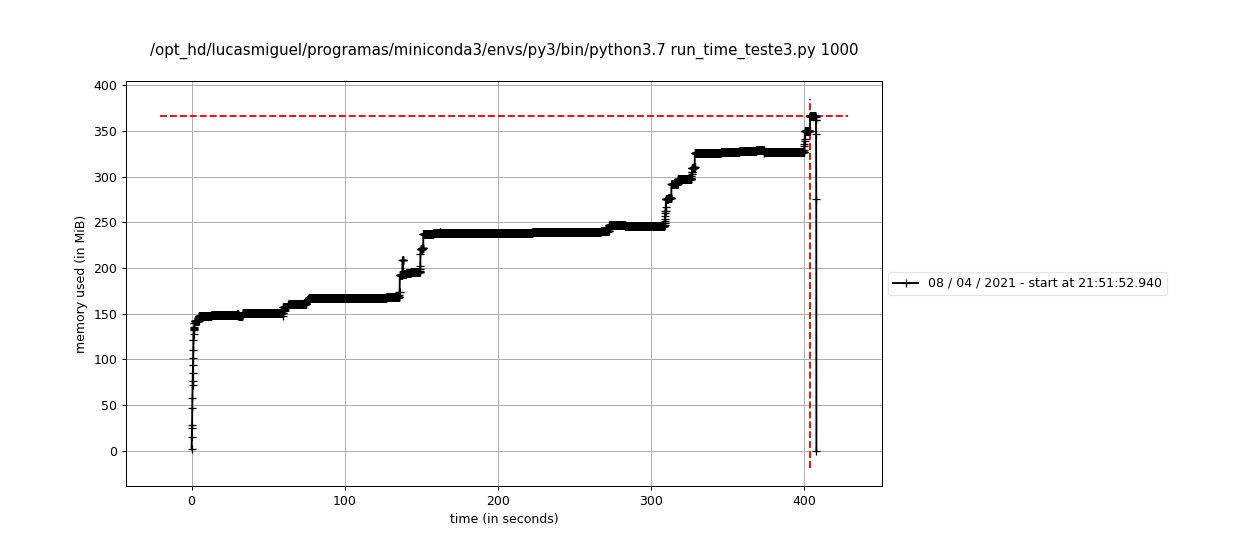

### plot_n1100.png

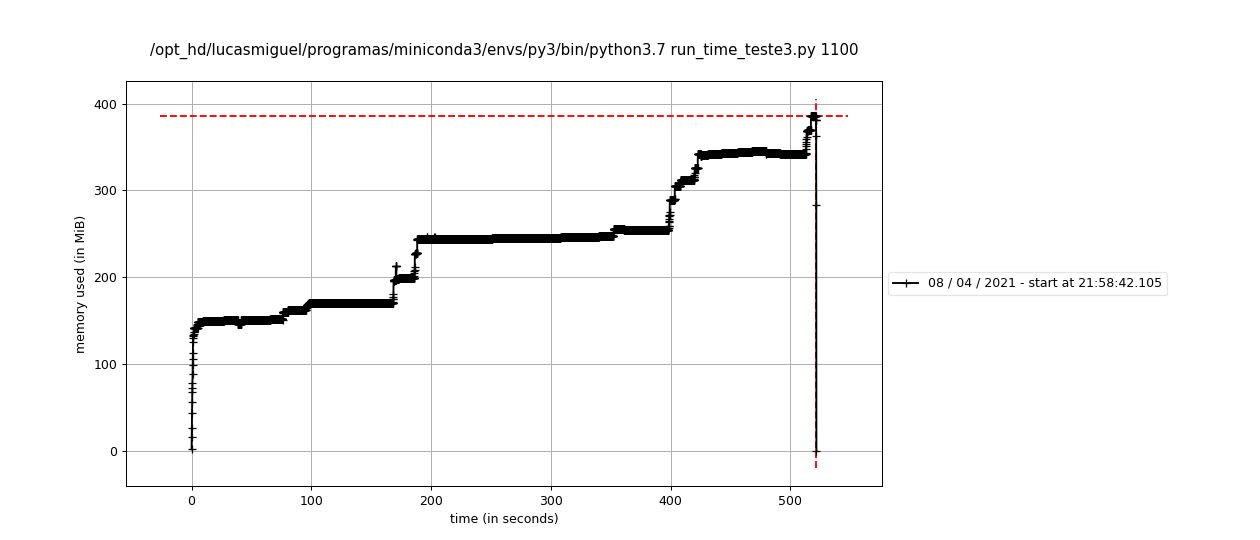

### plot_n1200.png

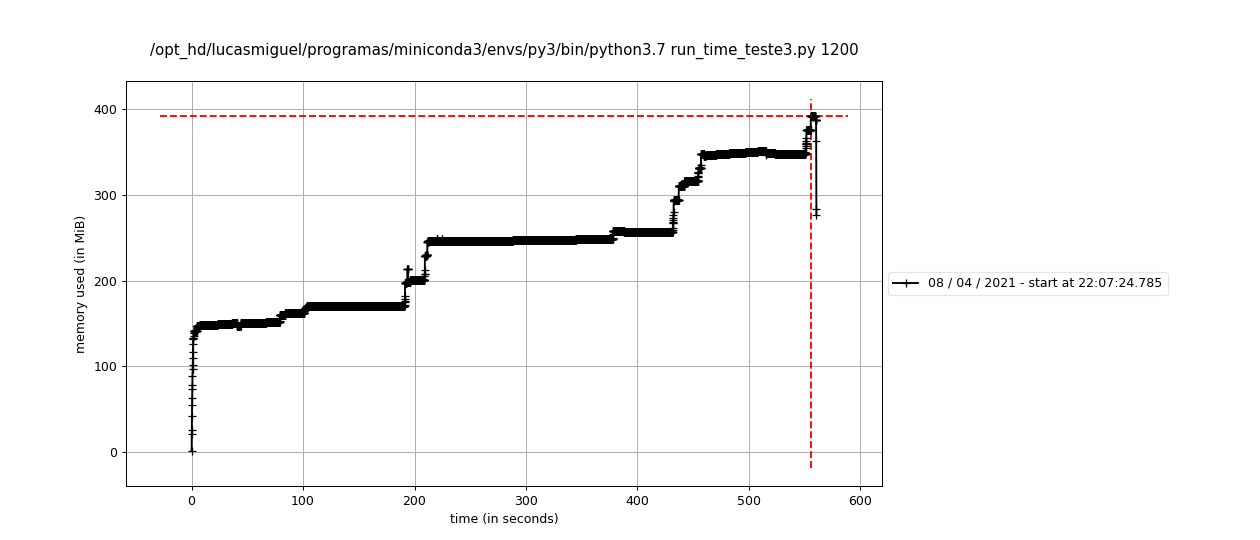

### plot_n1300.png

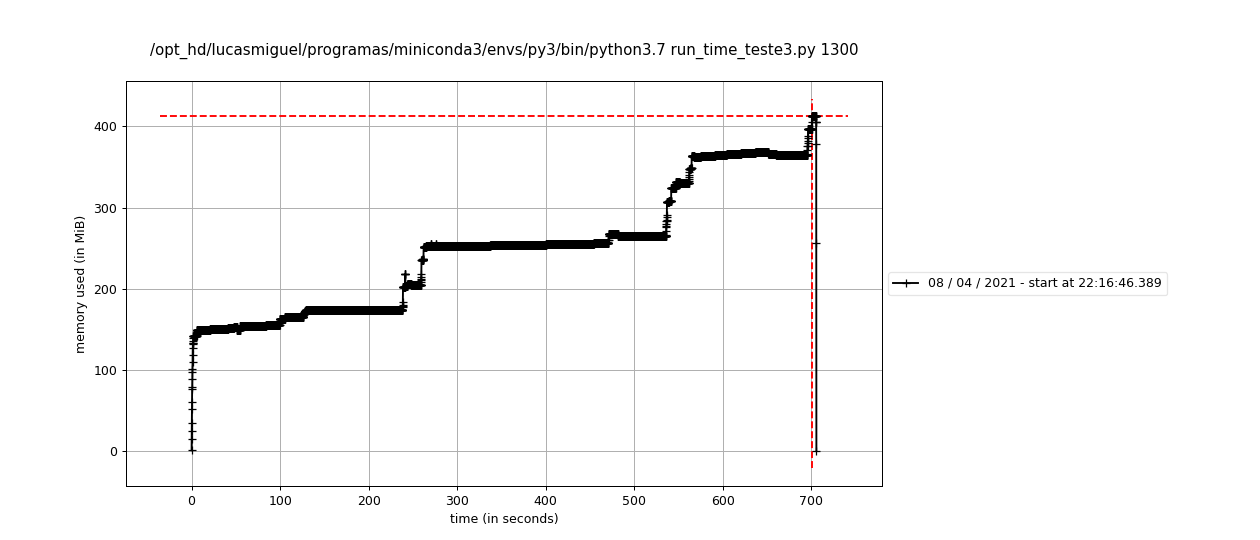

### plot_n1400.png

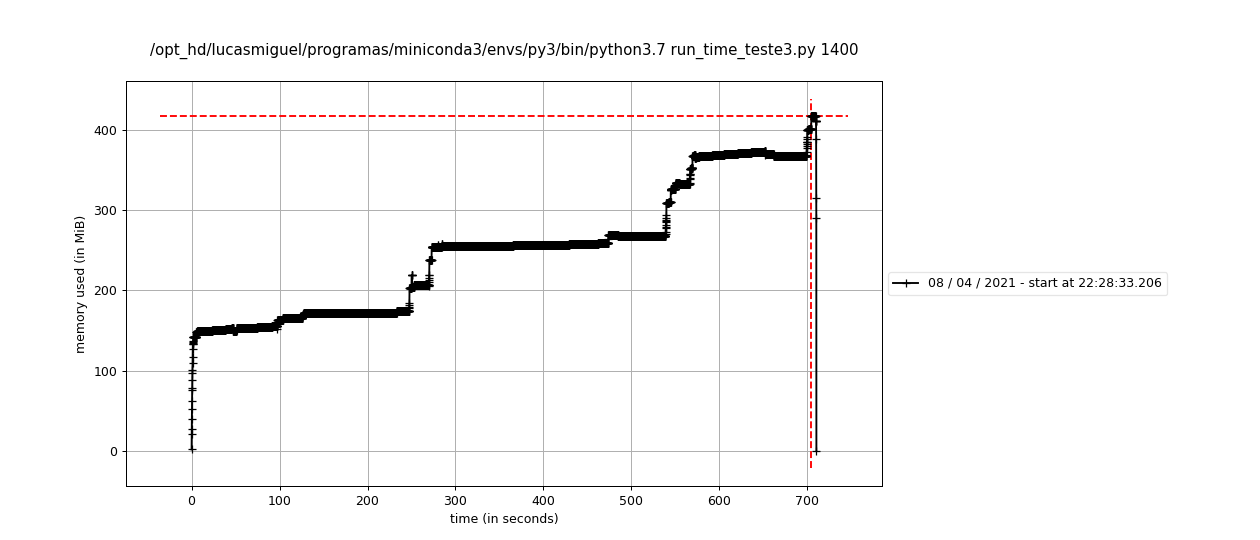

### plot_n1500.png

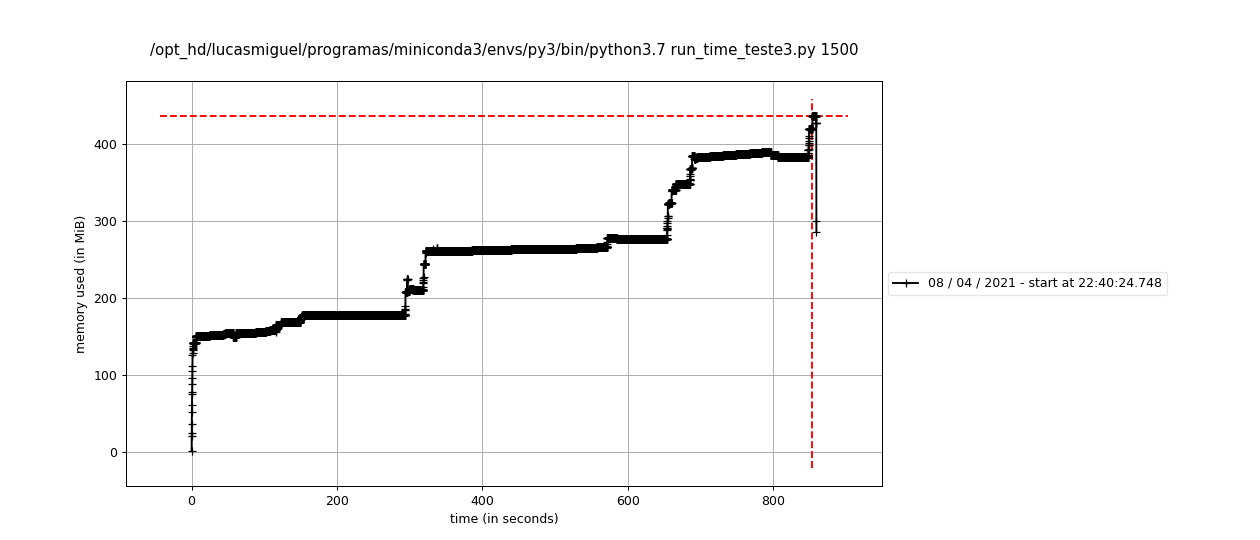

### plot_n1600.png

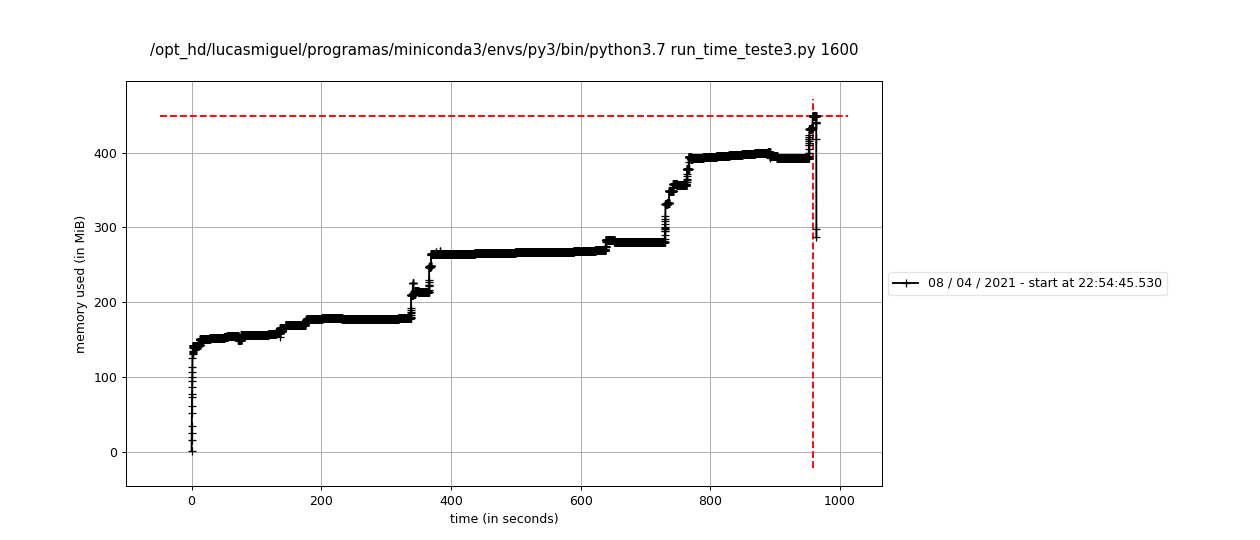

### plot_n1700.png

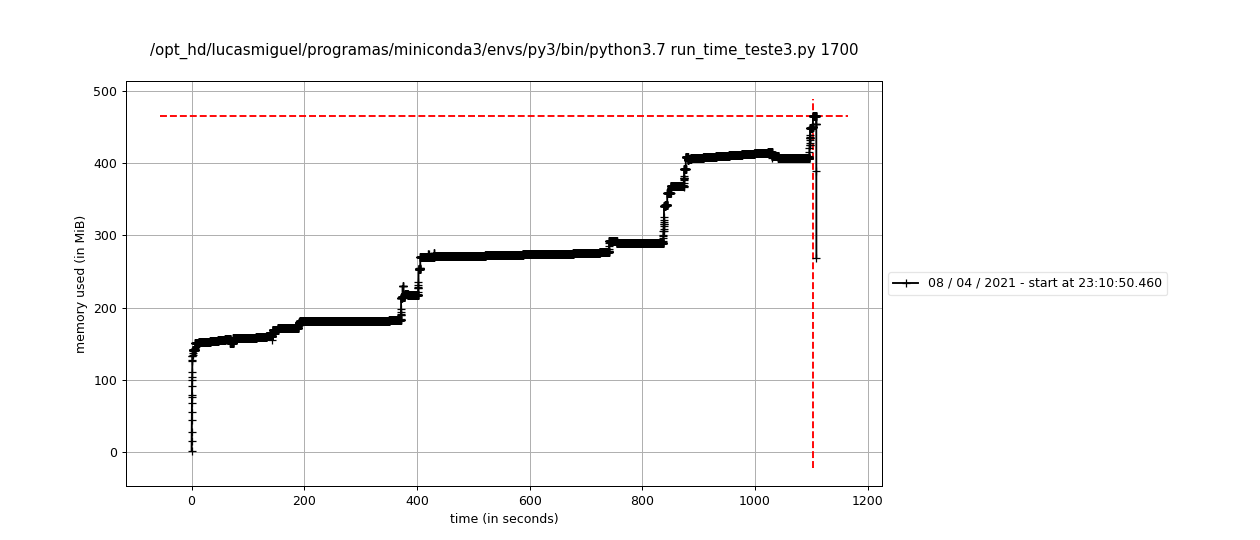

### plot_n1800.png

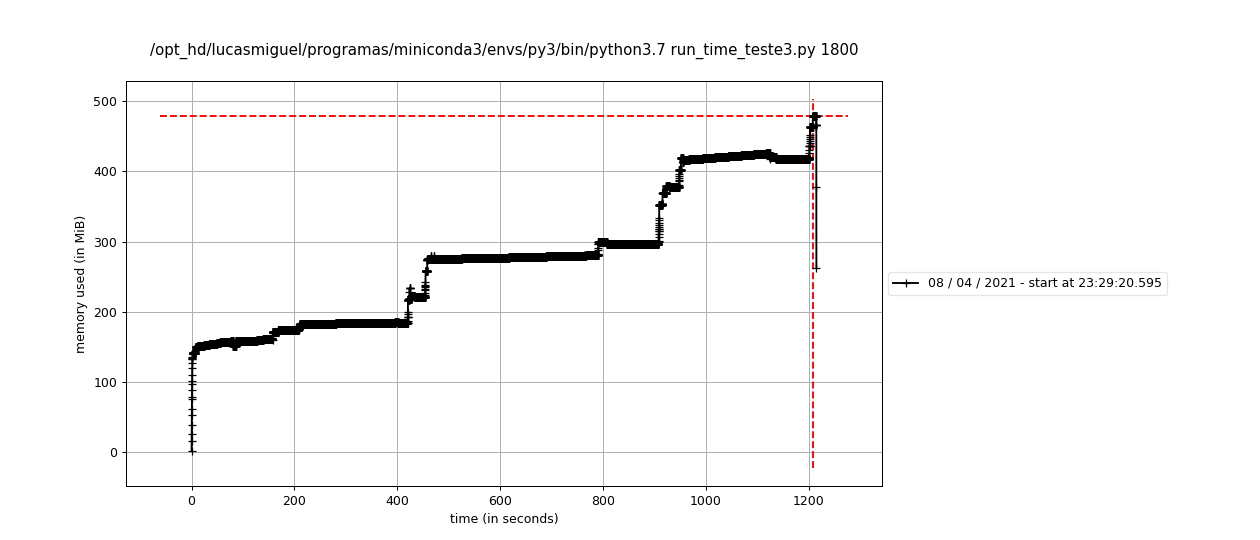

### plot_n1900.png

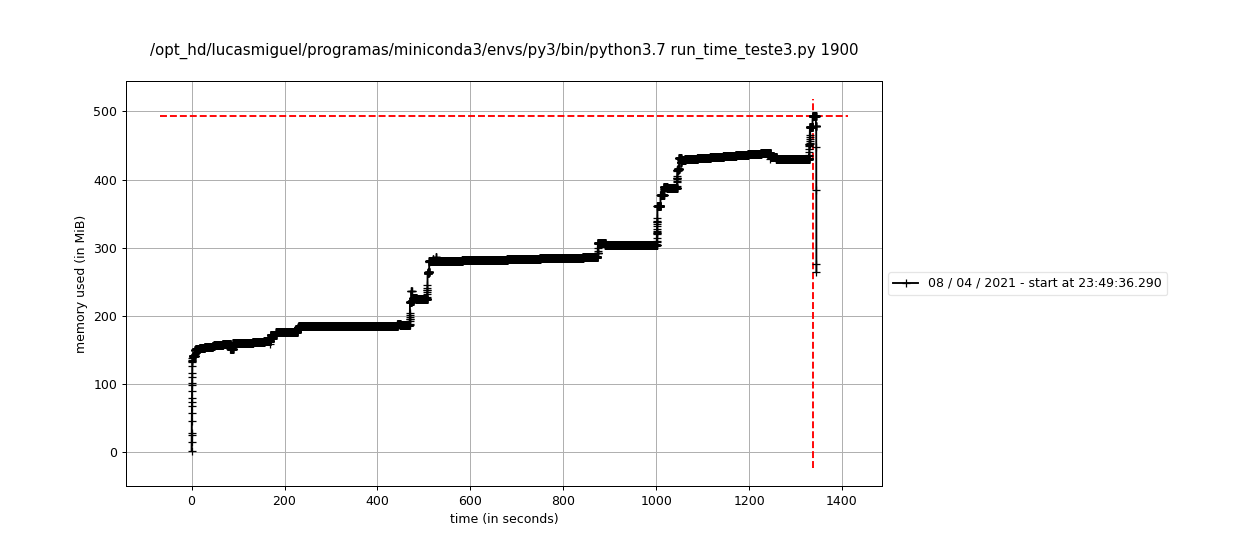

### plot_n2000.png

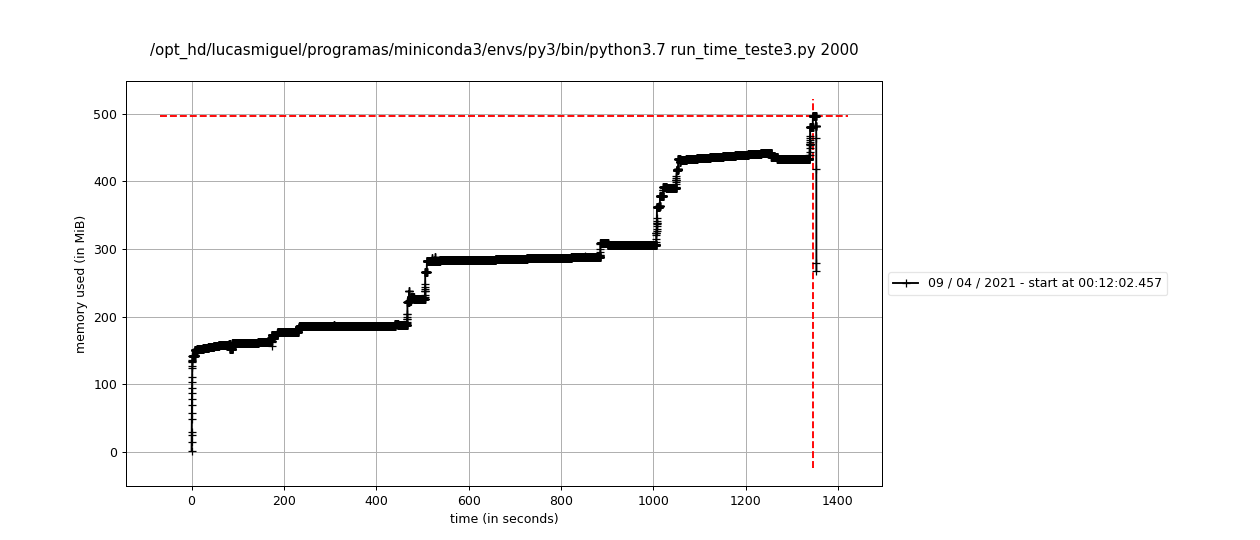
